# *Regulus* infers signed regulatory networks in few samples from regions and genes activities

**DOI:** 10.1101/2021.08.02.454721

**Authors:** Marine Louarn, Guillaume Collet, Ève Barré, Thierry Fest, Olivier Dameron, Anne Siegel, Fabrice Chatonnet

**Affiliations:** Univ Rennes, CNRS, Inria, IRISA - UMR 6074, F-35000 Rennes, France; UMR_S 1236, Université Rennes 1, INSERM, Etablissement Français du Sang, F-35000 Rennes, France; Laboratoire d’Hématologie, Pôle de Biologie, CHU de Rennes, F-35033 Rennes, France

**Keywords:** Activity patterns, Logical consistency, Few samples, Related cell types

## Abstract

**Motivation:** Transcriptional regulation is performed by transcription factors (TF) binding to DNA in context-dependent regulatory regions and determines the activation or inhibition of gene expression. Current methods of transcriptional regulatory networks inference, based on one or all of TF, regions and genes activity measurements require a large number of samples for ranking the candidate TF-gene regulation relations and rarely predict whether they are activations or inhibitions. We hypothesize that transcriptional regulatory networks can be inferred from fewer samples by (1) fully integrating information on TF binding, gene expression and regulatory regions accessibility, (2) reducing data complexity and (3) using biology-based logical constraints to determine the global consistency of the candidate TF-gene relations and qualify them as activations or inhibitions.

**Results:** We introduce *Regulus*, a method which computes TF-gene relations from gene expressions, regulatory region activities and TF binding sites data, together with the genomic locations of all entities. After aggregating gene expressions and region activities into patterns, data are integrated into a RDF endpoint. A dedicated SPARQL query retrieves all potential relations between expressed TF and genes involving active regulatory regions. These TF-region-gene relations are then filtered using a logical consistency check translated from biological knowledge, also allowing to qualify them as activation or inhibition. Regulus compares favorably to the closest network inference method, provides signed relations consistent with public databases and, when applied to biological data, identifies both known and potential new regulators. Altogether, Regulus is devoted to transcriptional network inference in settings where samples are scarce and cell populations are closely related. *Regulus* is available at https://gitlab.com/teamDyliss/regulus

## 1. Introduction

### Building patient-specific gene regulatory networks

Gene expression regulation (or transcriptional regulation) is a major field of investigation in life science. It allows a better understanding of major processes such as cell differentiation, cell identity and cell transformation (Garnis et al. 2004).

Chromatin accessibility has a major role in gene regulatory networks. Regulators are specialized proteins called transcription factors (TFs) which bind DNA in regulatory regions at specific binding sites. This results in activation or inhibition of target gene expressions. Regulatory regions are located in non-coding DNA (Smallwood and Ren 2013), and have to be in an accessible 3D conformation to allow regulation (Narlikar et al. 2002). Therefore, chromatin accessibility constrains transcription factor binding to DNA: a regulatory region in a “closed” conformation will prevent any potential regulation for a TF with a binding site inside it.

Gene regulation is extremely context-dependent (Duren et al. 2017; Sonawane et al. 2017; Ota et al. 2021): pathological processes such as cancer can disturb the transcriptional regulatory networks by modifying regulatory regions accessibility or location or by modifying TF fixation abilities (Khurana et al. 2016). Information about regulatory regions and TF expression is therefore essential to build reliable context-dependent gene regulatory networks and predict the effects of genome perturbations observed in pathological contexts, paving the way to personalized treatments. The majors challenges of studying the drivers of cellular evolution from patient-specific gene regulatory networks are (1) performing a subtle comparison of (2) closely related cell types (3) on a limited number of available samples.

### Use case: regulation driving B cells differentiation

As an illustration, let us consider the biological case study of differentiation of B cells into antibody producing cells. This process involves several transitions between closely related cell types, which are finely regulated by genetic networks (Willis and Nutt 2019). Their deregulation can lead to immunodeficiencies, autoimmune diseases and hematological malignancies. Few large scale studies have been performed to understand B cell normal and pathological differentiation. They all required a large number of samples to infer regulatory relations based on statistical analysis of TF and gene co-expression (Basso et al. 2005), or they only describe one type of B cells (Marbach et al. 2016). Other networks are only built with a limited set of regulators (Méndez and Mendoza 2016) or based on review of the literature. For example, (Willis and Nutt 2019) describes two main sets of opposed regulators: BACH2, PAX5 and BCL6 on the one side, inhibiting terminal differentiation and IRF4, PRDM1 and XBP1 on the other side, inducing it. However, a more complete characterization of B cells transcriptional regulatory networks is required to better understand the hijacking of the normal differentiation process by cancerous cells. To reach this goal, methods to infer regulatory networks from TF binding, gene expression and regulatory regions activity, obtained in few biologically-close samples and able to use the system dynamics to decipher the inhibition and activation roles of regulators, are needed.

### Issues with existing regulatory networks inference methods

The current transcriptional regulatory networks inference methods were reviewed for their ability to generate a reliable model of normal and pathological B cell differentiation, using the following criteria: (i) use of genes expression, TF binding and regulatory regions activity data, (ii) applicable to limited numbers of human samples, (iii) able to predict inhibitions and activations and (iv) reproducible and reusable on new datasets. The summary presented in Supplementary Table 1 shows that current methods are not applicable to limited numbers of closely related human samples, do not make a full use of all available information and rarely provide the activation or inhibition function of candidate TF-gene relations. These issues strongly limit the possibility to use such methods to better understand B cell differentiation.

### Regulatory Circuits and Semantic Web: proof-of-concept and limitations

The only reviewed method which uses regulatory landscape information and is applicable to human settings is the *Regulatory Circuits* project. This work is a major effort to merge large amounts of data from FANTOM5 (Andersson et al. 2014) to describe human regulatory networks (Marbach et al. 2016). The project resulted in 394 cell type-specific regulatory networks, in which TF-gene relations are weighted according to their strength in the considered cell type. As mentioned above, all the B cell subtypes of our case-study are included in a single network, preventing us to understand the differences between them and underlining a granularity issue.

In previous works (Louarn et al. 2019, 2022), we successfully integrated the *Regulatory Circuits* datasets into a unique structured graph in an attempt to analyze our case study with this pipeline. We used Semantic Web technologies, a generic data and knowledge integration framework (Berners-Lee and Hendler 2001; Berners-Lee et al. 2007) to generate a global RDF dataset that can be queried by dedicated SPARQL queries. This allowed to recompute the TF-gene relations published by the *Regulatory Circuits* project on all or a subset of the original datasets (including those related to B cells differentiation). Our study also highlighted that the *Regulatory Circuits* project design could not provide information about the activating or inhibiting role of regulating TFs. In addition, TF expression is not used as a selection criterion for relevant TF-gene relations. Finally, the *Regulatory Circuits* project methodology computes ranks for gene expressions and regulatory regions activities. This might be suitable for the large number of samples used in the original project, but not to describe changes between few closely-related cell types: it forces differences between very similar values, and it ignores the amplitude of activity changes.

### Objective

To address these issues, we introduce *Regulus*, an innovative gene regulatory networks inference tool to find the regulators of gene sets with similar expression dynamics and to qualify these TF-genes relations as activation or inhibition. *Regulus* has been developed to be stringent and to limit the space of the candidates TF-genes relations, highlighting the candidate relations which are the most likely to occur. *Regulus* is based on Semantic Web technologies, extending the methods introduced in (Louarn et al. 2019, 2022). Its main principles are to (i) take into account regulatory factors (TF and regions) activities, (ii) propose a discretization of the activities into patterns, (iii) produce signed networks inferred by a logical consistency step and (iv) be easily reusable and applicable to many datasets. By applying *Regulus* to published or original datasets, we show that it can describe regulatory networks with validated signed relations, it is able to identify known regulators of a specific biological process and it provides a list of new candidate regulators.

## 2. Results and Applications

### 2.1. The *Regulus* tool

We designed *Regulus*, a transcriptional regulatory networks inference tool dedicated to the analysis of few and biologically-close datasets. The tool relies on Semantic Web to integrate expression and epigenetic data. Figure 1 presents the different steps of the *Regulus* pipeline. It allows, from gene expression and regulatory regions activity tables, with their respective genomic coordinates, to infer consistent TF-gene relations.

**Figure 1:**
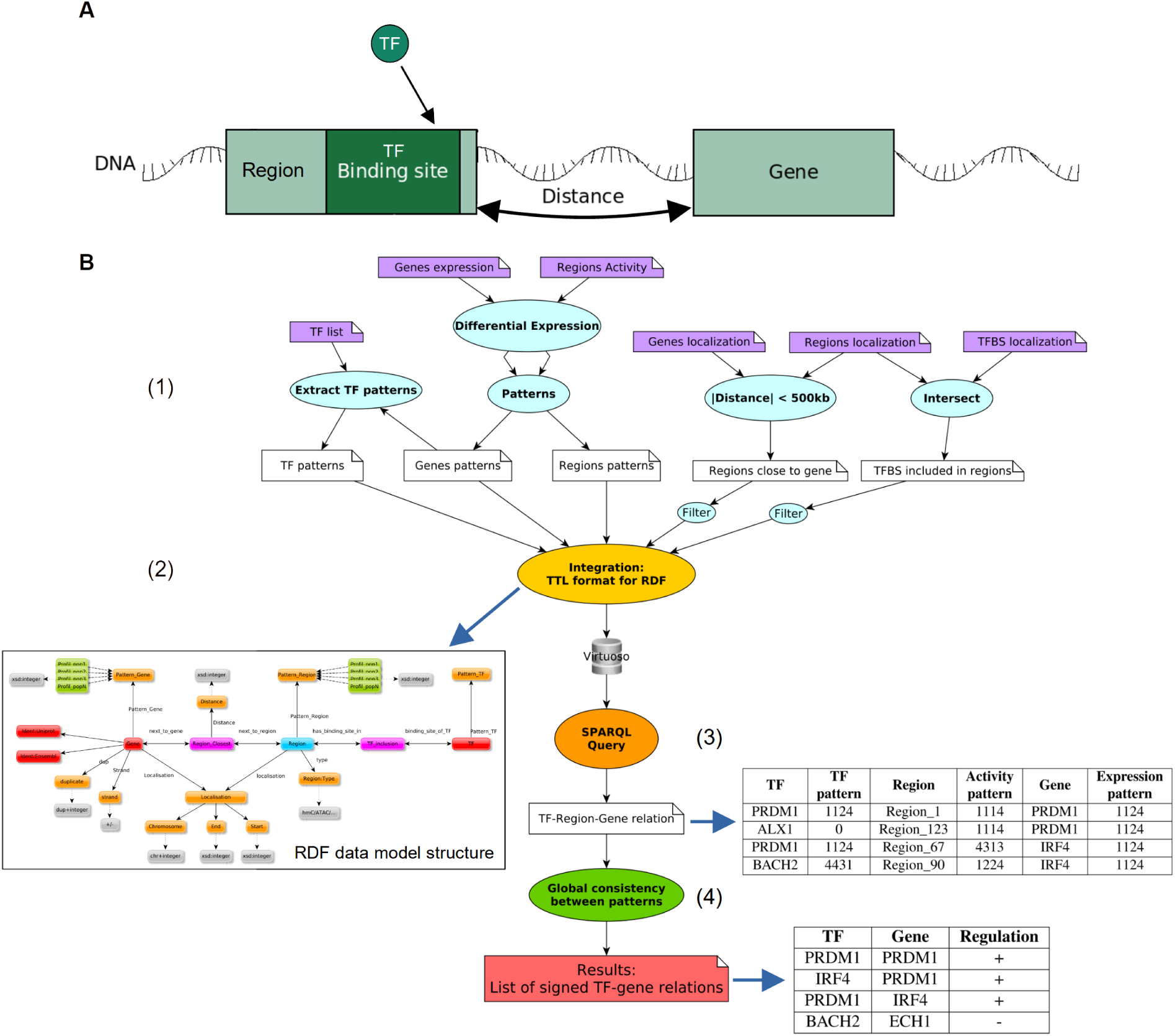
The *Regulus* pipeline. A: Scheme introducing the biological relationships between the different entities used to compute the transcriptional regulatory networks. B: Representation of the different steps of the pipeline. (1) Input data (purple boxes) are pre-processed (blue bubbles) according to the biology, expliciting the relations between the entities - inclusion of the TFBS in the regions (filtered to keep a single occurrence), distance between regions and genes (filtered to be less than 500 kb) - or reducing the data size and complexity by creating activity patterns for the different entities. (2) Integration and creation of a RDF graph formalizing the relations between all the entities (yellow bubble), generating a data model structure. (3) Dedicated SPARQL query to extract all the relations following the requirements introduced by Regulus (orange bubble) resulting in a global unsigned regulatory network expliciting relations as TF-Region-gene triples and their respective patterns. (4) Application of logical consistency constraints (green bubble) results in a signed and filtered regulatory network consisting in unique TF-gene relations (pink box).

#### 2.1.1. Pre-processing data for efficient integration: patterns as descriptions of data dynamics

The preprocessing steps (blue boxes and step (1) in the Figure 1B) consist in (i) finding relations between genes and regions by computing the distances between them, (ii) finding the inclusion of TF binding sites into regions, (iii) transforming individual entities (genes, TF and regulatory regions) activities (expressions and read densities) into patterns. The relevant entities for building a regulatory network are those of which the activity vary between the compared cell populations. Our network inference method is based on the common assumption that genes sharing a common expression dynamics are regulated by a common set of regulators (Yu et al. 2003). For each entity, *Regulus* therefore aggregates activity dynamics measured in *n* different cell populations into patterns, as a *n*-tuple. Each digit of this pattern represents a cell population and its value represents the relative activity of this entity in the considered cell population compared to all other populations, coded on four levels (1 to 4, see section 4.2 for details). Computed patterns therefore regroup entities which exhibit similar activity variations between cell populations regardless of their respective absolute activity levels.

#### 2.1.2. Data integration in a structure that can be browsed and queried

Pre-processed data are then integrated using Semantic Web technologies (yellow box and step (2) in Figure 1B). The data model structure after integration is illustrated in Figure 1B and Supplementary Fig 1, and can be seen as a representation of the interactions between the data, where the entities are linked between each other by explicit relationships. To retrieve TF-Gene relations, the strictly necessary entities are: genes, TF and regions with their respective patterns, as well as the reified entities Region_closest and TF_inclusion (see Section 4.3). Other entities or properties are kept to refine the results if needed.

Integrated data can then be interrogated to identify the candidate relations between entities that match a given set of rules: the regions must be at most at 500 kb of the gene, the TF must be expressed and have a binding site in the region (orange box and step (3) of Figure 1B). From the data structure, the corresponding SPARQL query is generated (Supplementary Fig 2): (1) starting from the node *Gene* with a specific expression pattern, (2) all *Regions* which are connected to the *Gene* by a *Region_closest* relationship are identified, (3) these *Regions* are associated to *TF* through *TF_Inclusion*, representing the presence of TF binding sites into regions. Along the way, the TF, regions and genes patterns are gathered, as they are required in the next step. The output of this step is a generegion-TF relations table (see (3) in Figure 1B)

#### 2.1.3. A logical consistency step for filtering and sign attribution

Query results must then be refined (green box and step (4) of Figure 1B) to prioritize TF-region-gene candidate relations consistent with the biological knowledge of how regulation works, and to infer a regulation sign (i.e. activation or inhibition) for these relations. To do this, a consistency principle is applied to the genes, TF and regulatory regions patterns.

This consistency rule is based on knowledge about the following biological principles for gene regulation: **(i)** the maximum effects on the gene expressions are obtained when the TF is at its highest expression level. **(ii)** The more accessible the region, the higher is the impact of the TF on the gene: for an activation this implies that if the TF is highly expressed so must be the gene, for an inhibition the higher the TF, the lower the gene’s expression. **(iii)** If the region looses its accessibility, the weight of the TF lowers: a higher TF expression level is needed to get a similar effect on gene expression. **(iv)** For fine grained tuning, two levels of confidence score are used to weight the consistency between the entities patterns and the relation sign.

These principles are transcribed into rules defining consistent sets of genes, TF and regions patterns and applied to score every single relation in each cell population. A global score for the relation is then calculated by adding these individual cell population scores and used to filter and sign the consistent relations (see Section 4.4). After using the consistency rule, the result is a quadruple between the gene, the neighboring regulatory region, the TF with a binding site in the region and the signed potential regulation on the gene expression. These relations can be merged into unique TF-gene relations when consistent relations involving the same TF and gene are found through different regions. The final output of the process is a signed TF-gene interaction table (see (4) in Figure 1B).

### 2.2. Application to *FANTOM5* data: validation of the basic principles

One aim of *Regulus* is to compute regulatory network on a limited number of samples and cell populations. *Regulus* was therefore run on four limited datasets extracted from *FANTOM5* and used in (Marbach et al. 2016), each containing four cell populations, chosen to be either of similar (comparable organ) or dissimilar (widely different localization) origin. Details about the chosen subsets are shown in Figure 2A. Data comprise 16,888 genes expressions, 43,012 regulatory region activities and 124,358,159 TF binding sites.

**Figure 2:**
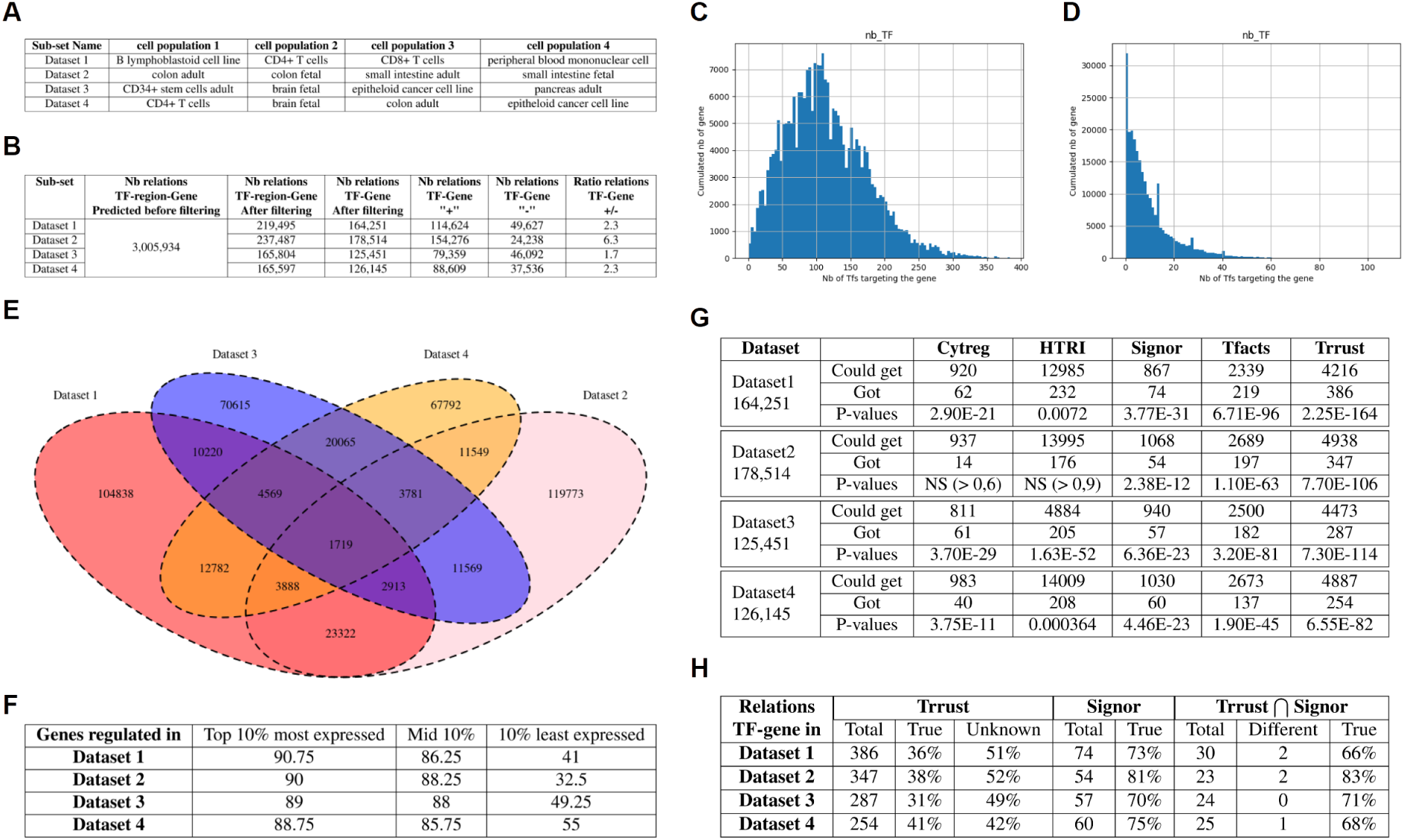
Applying *Regulus* to *FANTOM5* datasets produces reliable networks. A: Composition of the four datasets extracted from *FANTOM5*: two are composed of biologically-similar cell populations and two are composed of dissimilar cell populations. B: Number of relations inferred by *Regulus* for each of the datasets depicted in A. C-D: Distribution of the number of TFs regulating an expressed gene in *Regulus* networks before (C) and after (D) the filtering using consistency constraints. E: Overlap of the relations found in the regulatory networks corresponding to the four datasets of A. F: Percentage of genes from *Roadmap Epigenomics* RNA-seq datasets related to the cell populations found in *Regulus* inferred networks. Genes are separated in three categories according to expression levels: the top 10%, the middle 10% and the bottom 10%. G: For each dataset, number of relations which were present in each of the five database (“reachable”), number of these relations which were inferred by *Regulus* (“found”) and p-values of enrichment for existing relations in *Regulus* inferred networks, as assessed by binomial test. H: Relations found in Trrust and Signor and coherence of signs. True: percentage of relations found with the same sign as in the database, Unknown: relations non signed or signed + and - in the database. For the union of Trrust and Signor: different: number of relations signed differently in the two databases, True: percentage of relation signed the same as both databases.

#### 2.2.1. Regulus generates large networks which includes low expressed genes

##### Networks topology is refined by the consistency step

Figure 2B presents the number of relations obtained by *Regulus* for these four datasets. Just after the query and before running the consistency table, the same number of relations (3,005,934 TF-region-Gene or 1,869,854 TF-Gene) is generated for the four sets. This is due to the fact that exact same information about the TF binding sites and regions is used, leading to identical relations.

As seen in Figure 2B the consistency step allows to generate networks which are different in size and quality. They represent about 10% of the possible relations predicted by the query, underlining the filtering power of the consistency step. The consistency step modifies the network topology by modifying the distribution of the number of potential regulators per gene. As shown in Figure 2C-D, whereas in the total network genes can have up to 400 potential regulators with a mod of the distribution around 100, the consistency filtered networks have a more biologically realistic number of regulators per gene, ranging from 1 to 100 with a mod at 2 or Moreover, this filtering allows to prioritize relations which are relevant to the biological context, as seen in Figure 2E. Relations specific to each dataset are more numerous that common relations, these latter could be related to common biological processes necessary for basic cellular functions. This lower number of relations may be explained by the increased lack of consistency in TF-gene regulations across widely different cell populations.

Networks computed on dissimilar sets of cell populations are slightly smaller than the ones computed with similar subsets. This lower number of relations may be explained by the increased lack of consistency in TF-gene regulations across widely different cell populations. They also include a higher proportion of negative relations, which may be easier to identify when comparing strongly unrelated cell populations.

##### Inclusion of low-expressed genes

Gene inclusion in our networks is validated with the same *Roadmap Epigenomics* RNA-seq datasets used in (Marbach et al. 2016). As seen in Figure 2F, for the networks computed using *Regulus*, highly and medially expressed genes are recovered at high levels (> 90% for each category). Low-expressed genes are also significantly included in *Regulus* networks, with a retrieval rate ranging from 36.75% for similar cell types to 52% for dissimilar subsets.

#### 2.2.2. Inferred relations are validated by public data

##### Relations existence

To validate the relations inferred using *FANTOM5* data, several databases are queried: Trrust (Han et al. 2018), Signor (Licata et al. 2020), CytReg (Santoso et al. 2020), HTRIdb (Bovolenta et al. 2012) and TFacts (Essaghir et al. 2010)^1^. These databases contain manually curated TF-gene relations, found in cell types which may not be related to our datasets and only represent a fraction of all possible relations. Therefore their content is one or two levels of magnitude lower than the size of the networks inferred by *Regulus* and the number of common - or reachable - relations is even lower (Figure 2G). Altogether, our results show that *Regulus* inferred networks before the consistency step are highly enriched in relations described in each database (p-values ranging from 1.05e-59 to 2.25e-164). Moreover, for most cases (18 out of 20 comparisons), this is still true after applying the consistency step (p-values ranging from 0.0072 to 2.75e-288, Figure 2G).

##### Relations signs

Trrust and Signor both contain signed relations, but some can be unsigned or contradictory signed (within the same resource or between both), as evidence for activation or inhibition depends on the biological context. As shown in Figure 2H, the relations found in common between Trrust and Signor in our networks are signed the same way in both resources in most cases and in 72% signed the same way as in our networks. For the relations generated by *Regulus* and found at least in one of Signor or Trrust, the predicted sign is consistent with the databases in two third of the cases.

### 2.3. Application to B-cells identifies known and potential new key regulators

After having shown that *Regulus* performed well on closely related cell types with few samples, it is applied to new data about B cell differentiation.

#### 2.3.1. Biological context

After stimulation by a pathogen, naive B cells (NBC) differentiate into either memory B cells (MBC) or plasmablasts (PB). MBCs store some information about pathogen encounter and are able to differentiate faster and more efficiently if the same pathogen is present again. PB are effector cells and produce antibodies to inactivate and eliminate the foreign pathogens. Regulatory networks are required to explain the different abilities of NBC and MBC to differentiate into effector cells. For this study, four distinct populations are used: NBC, IgM^+^ MBC (MBC IgM), IgG^+^ MBC (MBC IgG) and PB. Gene expression data (RNA-seq, 26,734 genes) and chromatin accessibility data (ATAC-seq, 58,848 regions, used to determine regulatory regions activities) was generated. In this specific case the four populations are sequential: NBC is the first population and PB is the last, but MBC can be either a transitional state or a final one. Entities patterns will therefore have four digits, representing the relative entity activity in one B cell susbset in the following order: NBC, MBC IgM, MBC IgG and PB. Three main TF are highlighted in the bibliography at different steps of the differentiation: an inhibitor, BACH2, and two activators, IRF4 and PRDM1 (Willis and Nutt 2019), as shown in Figure 3A. We thus expect that *Regulus* infers these TF as main regulators in this system and in particular, that BACH2 is identified as an inhibitor of gene expression patterns increasing specifically in PB (such as 1114 or 1124, see Methods), but as an activator of decreasing gene expression patterns (4441, 4331 or 4431 for instance). PRDM1 and IRF4 are expected to have the opposite actions.

**Figure 3:**
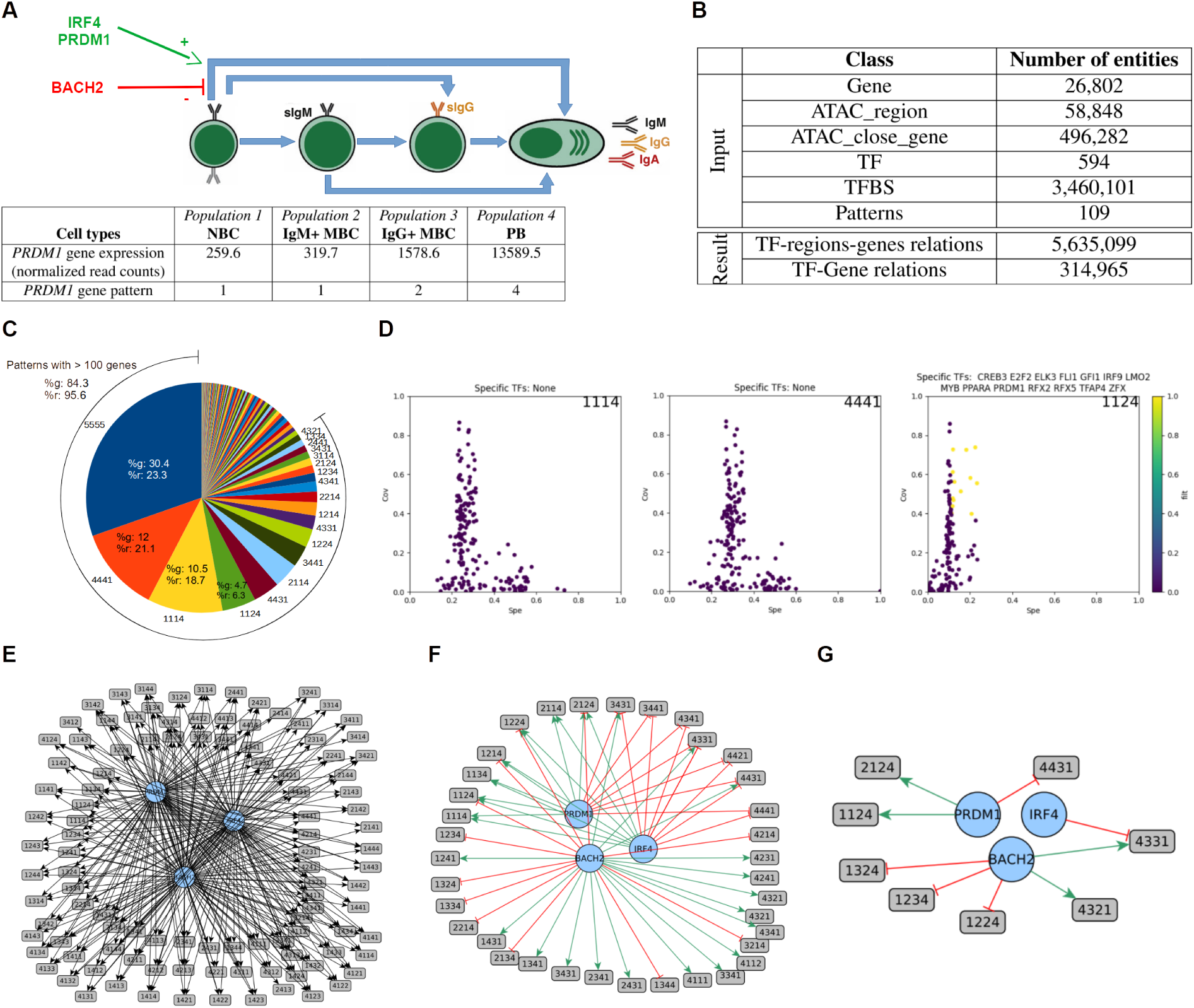
The B cell regulatory network computed with *Regulus*. A: Biological interaction between the cell types and main known regulators of the B cell differentiation process. Black arrows show the possible transitions between the different cell types. Cell types are ordered from the less differentiated (NBC) to the most differentiated (PB). BACH2 is a negative regulator of this process, whereas PRDM1 and IRF4 were described as positive regulators. Below the cells, an example of gene expression and its associated pattern is shown. The first pattern digit corresponds to NBC, the second to IgM+ MBC, the third to IgG+ MBC and the last to PB. NBC: naive B cells, MBC: memory B cells, PB: plasmablasts. B: Pie chart showing the number of genes per patterns of gene expression. The arc represents the 18 patterns with more than 100 genes, %g: percentage of total genes included in this pattern, %r: percentage of total relations targeting a gene included in this pattern. C: Table showing the main inputs and outputs of *Regulus* for the B cell differentiation regulatory network. D: Distribution of the coverage and specificity for all the TF targeting 1114 (left), 4441 (middle) and 1124 (right): yellow dots indicate the TFs passing the threshold. While being the largest patterns 1114 and 4441 do not have any key regulator with our filtering method. E-G: Graphs of interaction of the main known regulators and their targeted patterns before (E) and after (F) filtering the total relations with the consistency step and (G) after filtering with coverage and specificity: relations are consistent with the known roles of BACH2, IRF4 and PRDM1 during the PB differentiation.

#### 2.3.2. Patterns are indicative of expression dynamics

After differential expression analysis, 14,921 genes have a null or very low expression level in all samples and 3,591 genes are not differentially expressed in any comparison. These genes are respectively attributed the 0000 and 5555 expression patterns according to the convention described in Section 4.2.3. The remaining 8,222 genes are distributed over 107 patterns - among the 192 possible. As shown in Figure 3B and Supplementary Table 2, these patterns contain from 1 to 1,418 genes and 18 patterns are composed of more than 100 genes. It can be noticed that gene expression is not randomly distributed and that patterns indicate the main dynamics at work in our system: the most numerous patterns by far are 4441 and 1114. They show that many genes are down-regulated when either NBC or MBC are driven towards differentiation into PB - the B cell identity genes - whether another set of genes, necessary to mediate the PC differentiation and identity transition is induced (Willis and Nutt 2019). Interestingly, the third most numerous pattern is 1124 - containing the well known regulator PRDM1, which denotes the same dynamic as 1114, but with a small increase in IgG+ MBC, underlining their more advanced state of differentiation.

Among the genes, TF are treated as a specific entity since they are the potential regulators to be identified by *regulus*. In our dataset, 611 TF are retrieved from the 643 for which binding site information is available in *Regulatory Circuits*. 336 TF are unexpressed and hence will not impact the networks, leaving a set of 275 potential regulators in our system. TF expression patterns distribution is heavily unbalanced, since they are present in only 58 patterns. Moreover, 56 are constant genes, 63 have the 4441 pattern and 15 have the 3441 pattern, whereas only 11 and 13 TF are included in the 1114 and 1124 pattern, respectively. This suggests that most of the regulation supporting the NBC / MBC differentiation towards PC is due to TFs which expression is extinct in plasma cell and is mainly a release of inhibitions allowing the PC fate driving regulators and genes to be expressed.

Finally, regulatory regions are divided over 110 patterns with a distribution which is highly correlated to the gene expression patterns (*r*^2^ = 0.89 including all patterns, *r*^2^ = 0.68 when the constant patterns are removed). As for the genes, the 4441, 3441, 1114 and 1124 patterns are the most numerous, underlining that according to the biological rules of gene regulation, the above-mentioned gene expression inhibitions and inductions are sustained by concordant changes in regulatory regions accessibility.

In summary, pre-processing the data into patterns provides some filtering power, allows the user to concentrate on patterns relevant to the biological context and gives a first glance of the dynamics at work in B cell differentiation.

#### 2.3.3. Potential relations need to be consistency-filtered

Pre-processed data are then integrated to create the data structure and the relation graph and queried. *Regulus* inputs and outputs are shown in Figure 3C. The pipeline generates 5,635,099 TF-regions-genes relations, which resulted in 314,965 unique TF-gene signed relations, once filtered with the consistency table and merged to regroup TF-gene relations that occur through different regions. Of those relations 173,717 are signed as activation (+) and 141,248 as inhibition (−). As described in Figure 3B and Supplementary Table 2, the most numerous gene expression patterns (except the constant one) have a tendency to aggregate a larger proportion of relations compared to the percentage of total genes they include. As shown in Section 2.2, in Figure 3C and by comparing Figure 3E and Figure 3F, the consistency steps refines the resulting network to the potentially most relevant relations.

#### 2.3.4. Key regulators can be identified by coverage and specificity filters

As a further refinement to identify the most biologically relevant TF-gene (or TF-pattern) relations, the stand-alone tool *ClassFactorY* is provided, allowing to apply a coverage and specificity filter and to provide an annotation-based score (see 4.5.1). After using the specificity and coverage filter (using the last quartile as a threshold for both), no specific TF is highlighted for 25 patterns, a single potential regulator is found for 16 patterns and 10 or more regulators of interest are identified for 35 patterns. Surprisingly, for the two non-constant most populated patterns (1114 and 4441), no TF could be singularized by the coverage / specificity filter (Figure 3D). Out of the 238 TF found in the resulting relations, 116 do not pass the thresholds for any pattern, 23 regulate only 1 pattern and 19 are associated with 10 patterns or more. The combination of both parameters also allows us to filter out some highly ubiquitous TF, such as SP4.

Among the 121 TF passing the threshold, many have already been described as implicated in B cell differentiation, including the main known regulators PRDM1, IRF4 and BACH2 (Willis and Nutt 2019). At the pattern level (Figure 3G), IRF4 is found as an inhibitor of 4331, PRDM1 is an activator of 1124 & 2124 (strong expression specifically in PB) and an inhibitor of 4431 (low expression in PB). BACH2 is identified as an activator of 4321 & 4331 (decreasing expression during differentiation) and an inhibitor of 1224 & 1234 & 1324 (increasing expression during differentiation). *Regulus* results are therefore in agreement with the literature and with our expectations (see 2.3.1). Complete metrics about TF-gene relations based on the number of genes in each pattern and describing the number of relations, of potential regulators before and after the coverage / specificity filter and some biological interpretation are available in Supplementary Table 2. The refining power of both the consistency table and the coverage / specificity filter is appreciated by comparing the resulting networks in Figure 3E-G.

#### 2.3.5. Regulators annotation as a decision helping step

*ClassFactorY* also includes an annotation based score (see 4.5), based on GO terms, Pubmed citations and MesSH terms allowing the validation of already known TFs, relevant in the biological context, and providing a list of potential TFs of interest for further biological investigation. Out of the 121 TFs identified in the previous step, 64 are annotated by the GO term “cellular developmental process”, 27 by “immune system process”, 13 by “lymphocyte activation”, 3 (IRF8, LEF1, YY1) by “B cell activation”, 2 (IRF8, YY1) by “B cell differentiation” and none by “plasma cell differentiation”. The number of citation in Pubmed for the found TFs ranges from 164,841 (MAX) to 2 (ZNF75A).

Citations involving MeSH terms relative to the specific context of the B cell differentiation and each of the TF are counted: 93 TFs are cited, among which IRF4, STAT3, MYC, PRDM1 & PAX5 being the five with the most citations (208 to 386). A second query is done with a more global context relative to B cells in general: 104 TFs are annotated; PRDM1, PAX5, STAT3, MYC & MAX being the 5 with the most citations (402 to 2573). A null score is attributed to six TFs (KLF16, FOXJ3, TFAP4, TGIF1, ZNF219 and ZNF75A), meaning that although they have been selected by the coverage / specificity filter, their potential role in B cell has not been studied and may be worth investigating.

### 2.4. Regulus compares favorably to Regulatory Circuits

After validating our pipeline, it is compared to the closest method of network inference in terms of dataset size and nature of the genomic features, *Regulatory Circuits*.

#### 2.4.1. Workflows comparison

Figure 4 presents the main steps of the *Regulatory Circuits* pipeline (Figure 4A) and compares them to those of *Regulus* (Figure 4B). Both take as input the regions and genes activities and localizations, as well as TF binding sites coordinates. Both also share similar pre-processing steps like computing the distance between regions and genes or finding the TF binding sites occurrences in the regulatory regions. The main differences are: (1) *Regulus* uses the activity of the TF (extracted from its coding gene activity) which is not taken into account by *Regulatory Circuits* workflow; (2) *Regulatory Circuits* uses a composite score in which each component must be strictly positive, and is taken as a maximum when several concordant relations exist. This approach brings a bias towards activation relations and favors highly expressed genes. On the contrary, *Regulus* checks the global consistency between the different activities of the relation entities over all cell populations to produce signed networks; (3) *Regulatory Circuits* gives cell population-specific networks whereas *Regulus* outputs networks by patterns, adding dynamics to the network.

**Figure 4:**
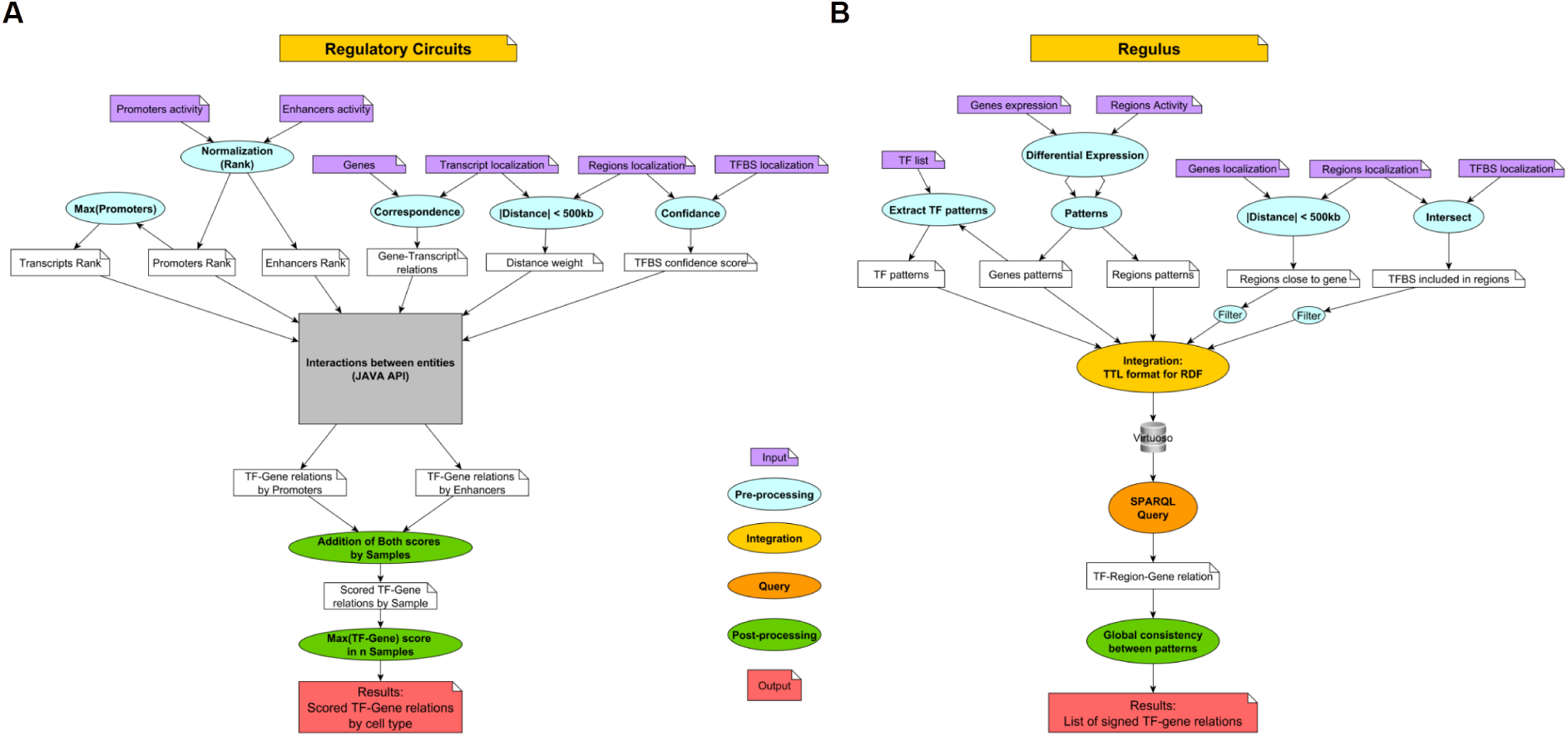
Comparison of the *Regulatory Circuits* (A) and *Regulus* (B) workflows: the pre-processing (in blue) steps are similar: normalization of the expressions, limitation of the distance between region and genes and inclusion of the TFBS in the regions. The main differences are: *Regulatory Circuits* uses a rank normalization of entities activities while *Regulus* uses a clusterization by pattern. While the construction of the regulatory network use different technologies (gray, yellow and orange) they follow similar principles: finding the biological relations between the entities and retrieving the weight (A) or patterns (B) along the relations. The main differences are in the post-processing (green): *Regulatory Circuits* gives a score for each relation in a sample, filters out the null scores and computes the cell populations networks, while *Regulus* computes a regulatory network, filters it with a consistency rule and signs every interaction.

#### 2.4.2. Networks and outputs comparison

##### Relations found in reference databases

The same databases as in 2.2.2 are used to evaluate the results of *Regulatory Circuits*. Overall, *Regulatory Circuits* produces similar enrichment of relations found in databases as *Regulus*, with significant enrichment obtained in 19 out of 20 cases (p-values ranging from 4.6e-3 to 1.7e-290).

##### Networks topology

For the *FANTOM5* datasets used in 2.2, *Regulatory Circuits* number of potential TF-genes relations (2,060,960) is calculated based on the supplementary data files describing entities (TF, promoter, enhancer, transcript) relations, while ignoring the scores (see (Marbach et al. 2016)). All the TF-gene relations found by *Regulus* are included in the potential *Regulatory Circuits* relations, meaning that our method does not create irrelevant relations.

However, *Regulatory Circuits* provides very large networks, comprising from 407,056 to 1,796,098 unsigned relations for a single cell population; whereas *Regulus* signs and refines these relations to less than 180,000 for sets of four different cell populations, adding an interesting filtering power. Interestingly, it shows the same “number of TF per gene” distribution than the non consistency-filtered *Regulus* output (see Supplementary Figure 4A-C).

##### Inclusion of genes based on their expression level

When using a similar validation of output genes using Roadmap Epigenomics RNA-seq datasets as in 2.2, *Regulatory Circuits* is well able to retrieve highly or moderately expressed genes in its networks. However, lowly expressed genes are poorly incorporated, in agreement with their methodology favoring inductive relations and high activities (see above). On the contrary, *Regulus* manages to retrieve similar percentage of highly and moderately expressed genes but higher percentage of lowly expressed ones (Figure 2F and Supplementary Figure 4D-E).

## 3. Discussion

In this article we present a new design for regulatory network inference. *Regulus* addresses some of the methodological issues presented in Introduction: (1) the under-exploitation of the regulatory context, (2) data reduction and structure, (3) the lack of functionality qualifier for interactions (activation of inhibition) and (4) pipeline availability for reuse and reproducibility. To solve theses issues (1) the TFs expression and regulatory regions activities are used by the pipeline, (2) genes or regions of similar activity trend are clustered under the same pattern and (3) data is integrated in a Semantic Web technologies based structure, that can be browsed and queried. A global network based on all the samples is computed and the consistency of the relations with the biological background is checked, allowing to sign the relations and to not discriminate the inhibition. (4) An automated version of *Regulus* is provided to facilitate its reuse. Altogether, *Regulus* compares favorably to the only similar method incorporating knowledge about regulatory regions, *Regulatory Circuits*.

The solution chosen to cluster genes of similar expression direction in cell populations is to group them by patterns. Describing the expression or activities as patterns allows the user to concentrate on patterns which have a biological meaning in the given experimental setting. Cell types are not individualized by these patterns, but biologists are often more interested in dynamic changes between cellular states than by a complete record of regulators active in a fixed cell population. Even if they are interested in such characterization, it is possible to look at regulators for the pattern where expression is at its highest only in the cell population of interest. A limit of this approach might be the over-sampling of the patterns as some patterns were very poorly populated: 33 with less that 10 genes including 18 with less than 5 genes on the B cells data. Those patterns can be grouped with patterns of similar direction or removed from the analysis, since they may bias the coverage and specificity filter. Indeed small patterns are easily covered and may introduce high specificity percentages. Another solution would have been to use co-expression analysis for example using the WGCNA R (Langfelder and Horvath 2008) package. Unfortunately, after testing, it gave poor results with our limited number of datasets.

Our work shows the added value of integrating large and heterogeneous data from biological experiments when inferring regulatory networks. Even on closely-related cell types, such as the B cells subsets, subtle differences can be identified owing to the use of patterns and the consistency step. Semantic Web Technologies allow for an easy identification of relations. Once the data structure is obtained, it can be queried to answer any specific question. As it is based on unique identifiers and self-structuring, it also reduces the risks of introducing false relations, which may happen when manipulating text files with command line tools. This confirms that Semantic Web technologies, which were instrumental in the expansion of the Linked Open Data initiative (Bizer et al. 2009), and in particular in life science data integration, (Kamdar et al. 2019; Kamdar and Musen 2021), are also a suitable framework for supporting gene-regulatory network inference in the context of personalized medicine.

The *ClassFactorY* tool is provided to identify the key regulators by introducing a filter of coverage and specificity, and by adding biological context annotations. From the 237 potential TF involved in B cell networks, the coverage and specificity reduces this number to 121, which is still too much to perform experimental validation. Annotations can thereafter be used on these “short-listed” TF to (1) validate known regulators and (2) identify potential new regulators which have not been described in the context of interest. A perspective is to reduce the number of candidate regulators by using constraint programming to determine the smallest group of TF able to regulate the biggest part of a gene pattern.

Finally, the combination of *Regulus* and *ClassFactorY* is able to retrieve the main regulators of the B cell differentiation process, such as PRDM1, IRF4, BACH2 and PAX5 and to pinpoint them with a high annotation score. On the other hand, six new potential TFs impacting this process, identified through high coverage and specificity coupled to a null annotation score, will need to be further investigated, showing the power and interest of our tool.

## 4. Methods

### 4.1. Main characteristics of *Regulus*

As inputs, *Regulus* requires: (i) a list of genes with their expression in a selected number of cell types, (ii) a list of selected regulatory regions with their activity as two text files, (iii) genomic locations of genes, regions and TF binding sites as bed files. The latter can be provided by the user (ChIP-seq data for a specific TF for example), but the genome-wide TF binding sites coordinates from *Regulatory Circuits* (Marbach et al. 2016), containing curated binding sites for 643 TF, are provided. All genomic coordinates are given according to the *hg19* human reference genome. *Regulus* outputs a list of candidate signed TF-genes relations with patterns indications for all entities, that can be explored to identify new regulators.

### 4.2. Pre-processing in *Regulus*

#### 4.2.1. Neighborhood relationship

Genomic coordinates are used to compute distances between regions and genes with a custom *python* script. The distance is calculated between the two closest extremities of the entities, regardless of their respective position. All distances are filtered at a max threshold of 500 kb and set to 0 for overlapping entities.

#### 4.2.2. Finding TF binding sites in our regions

*Regulatory Circuits* (Marbach et al. 2016) data on TF localization across the genome are used, as they contain reliable and extensive information. The *Bedtools intersect* (Quinlan 2014) tool is first used to identify all the TF binding sites included into a set of regions. For a given TF, only the occurrence of at least one binding site per region is kept, producing binary relations between TFs and regions.

#### 4.2.3. Gene Expression and Region Density Patterns

Entities (genes, TF or regions) with differential activities (expression or accessibility) are defined by the user as input and grouped into patterns, according to activity dynamics. This discretization is performed independently for each entity, as activity levels may greatly differ between entities (Marbach et al. 2016). A pattern is a *n*-tuple with *n* equal to the number of compared cell populations. First, the mean per population based on samples (one population = several samples) activities (normalized read counts, reads densities…) is calculated and log-transformed. The interval between the maximal and the minimal of these average values is divided into four equivalent intervals, providing a scale from 1 (minimum) to 4 (maximum). Each averaged expression value, and therefore each cell population, then gets an attribute from 1 to 4 corresponding to the interval where this value belongs.

As an example Figure 5 shows the pattern attribution of an entity activity over four cell populations, in the form of a quadruple of integers, where the entity is a gene and its activity measure is its averaged expression. Figure 5A shows the gene expression (in normalized read count per million) and the attributed pattern value for each cell population. Figure 5B illustrates the pattern generation: the highest activity level (log10 transformed) is set to 4, the lowest is set to 1 and the space between both is divided in four equal intervals, numbered from 1 to 4. Each cell population gets the interval number corresponding to its activity level as a pattern value, and these values are aggregated in a quadruple, in this case leading to the pattern 1124.

**Figure 5:**
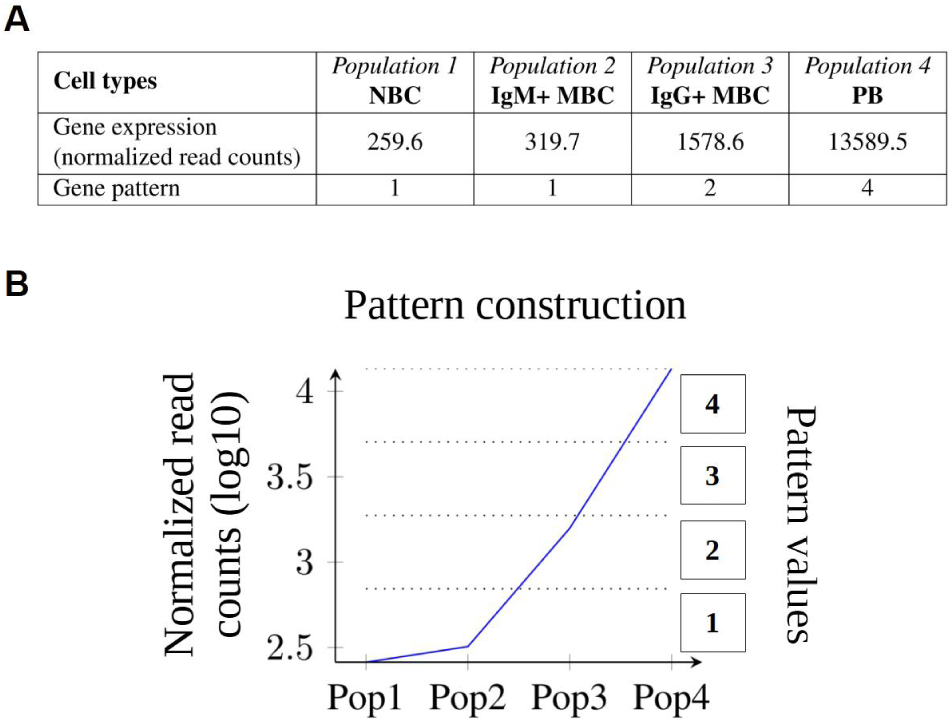
Example of a gene expression pattern construction. A: This gene expression is characterized by two low expression values followed by an intermediate and a very high value. B: This expression is modeled by the 1124 pattern after log transformation and binning of the difference between minimal and maximal values in four equal intervals.

Finally, entities with no or low activity (as defined in differential expression analysis for genes, for example) in all samples are granted a profile with only zeroes as values and are removed from the implementation, since they do not bear any relevant biological information. A profile with *5* for all cell populations is attributed to entities with constant activity. The TF patterns are those of their respective coding genes.

### 4.3. Data graph for the integration and query in *Regulus*

Pre-processed data are transformed from tabulated to TTL format by a custom *python* script, allowing to introduce the inclusion relation between TF and regions, or the distance relation between regions and genes>Then, *Regulus* generates a structured RDF graph of data that can further be queried with Semantic Web technologies (Supplementary Fig 1). The construction of the RDF dataset requires the introduction of unique identifiers and of reified entities (Hernández et al. 2015; Nguyen et al. 2014) to describe some relations. (1) <u>Identifiers</u> For the regions, a unique identifier is designed after the type of region (i.e. ATAC_; and Region_ in the text) followed by the row number at which they appear in the region localization file - this ensures that a same number is never used twice for different entities. For genes and TF, their HGNC Gene Symbols are kept as identifiers. (2) <u>Reified entities</u> As RDF does not allow relations to bear a score, relations of distance between genes and regions and of TF binding into regions are both inserted by using reified relations. This method generates new entities with devoted identifiers and bearing either the distance value or the inclusion. (3) <u>Additional information</u> References to Uniprot and Ensembl are added for the genes to link our data with public databases. The localization information for all entities is also kept, as a potential filtering parameter.

To find the potential regulators of a given set of genes, *Regulus* uses a SPARQL query (Supplementary Fig 2) on the previously generated data graph to extract all TF-Region-Genes triples and their related patterns.

### 4.4. Consistency table to assign roles to relations in *Regulus*

*Regulus* relies on a principle of consistency between genomic landscape, genes and TF expressions to decide if a relation is susceptible to exist and to predict if this relation will act as an inhibition or an activation. Consistency constraints are based on biological principles, as detailed in section 2.1.3 and transformed into rules applied to each value of all entity patterns, determining a consistency score for each studied cell population. These scores have two confidence levels and are stored into tables shown in Supplementary Figure3. Scores are assigned to each cell population by choosing a sign and looking first at the region pattern value, which determines the assignment table containing the scores. Then the TF (rows) and gene (columns) pattern values are used to determine the score, which is at the intersection of the corresponding values.

Figure 6 presents a graphical representation of the consistency principles application. First (top of Figure 6), region, TF and gene entity patterns are retrieved. In this example, the chromatin is closed and the TF is lowly expressed in the first population (Figure 6 middle line). To decide if the relation is more likely to exist as an activation or inhibition in this cell population, *Regulus* looks into the two tables (activation, middle left and inhibition, middle right) corresponding to a region pattern value of 1. The TF and gene each have a pattern value of 1 in this first cell population, as indicated by the light blue and yellow boxes in the central table of Figure 6. Therefore the assigned score for this population for an activation is +2 and 0 for an inhibition, as shown by the intersection of the corresponding lines and columns in the score assignment tables. For the last population the chromatin is highly accessible and both the TF and gene are highly expressed (pattern values of 4, bottom part of Figure 6). In this case, tables at the bottom are used (according to a change in the region pattern value) and the scores are determined by the intersection of the dark blue and olive boxes, being again +2 for an activation and 0 for an inhibition. These steps are also performed for cell populations 2 and 3 (not shown) and the individual scores are added to get a global consistency score (end column of the central table in Figure 6), either for activation (green colored cell) or inhibition (red colored cell), which can be filtered by a threshold. For example this threshold is fixed to 7 for a 4 digit pattern, allowing at most one individual cell population score to be of lower confidence.

**Figure 6:**
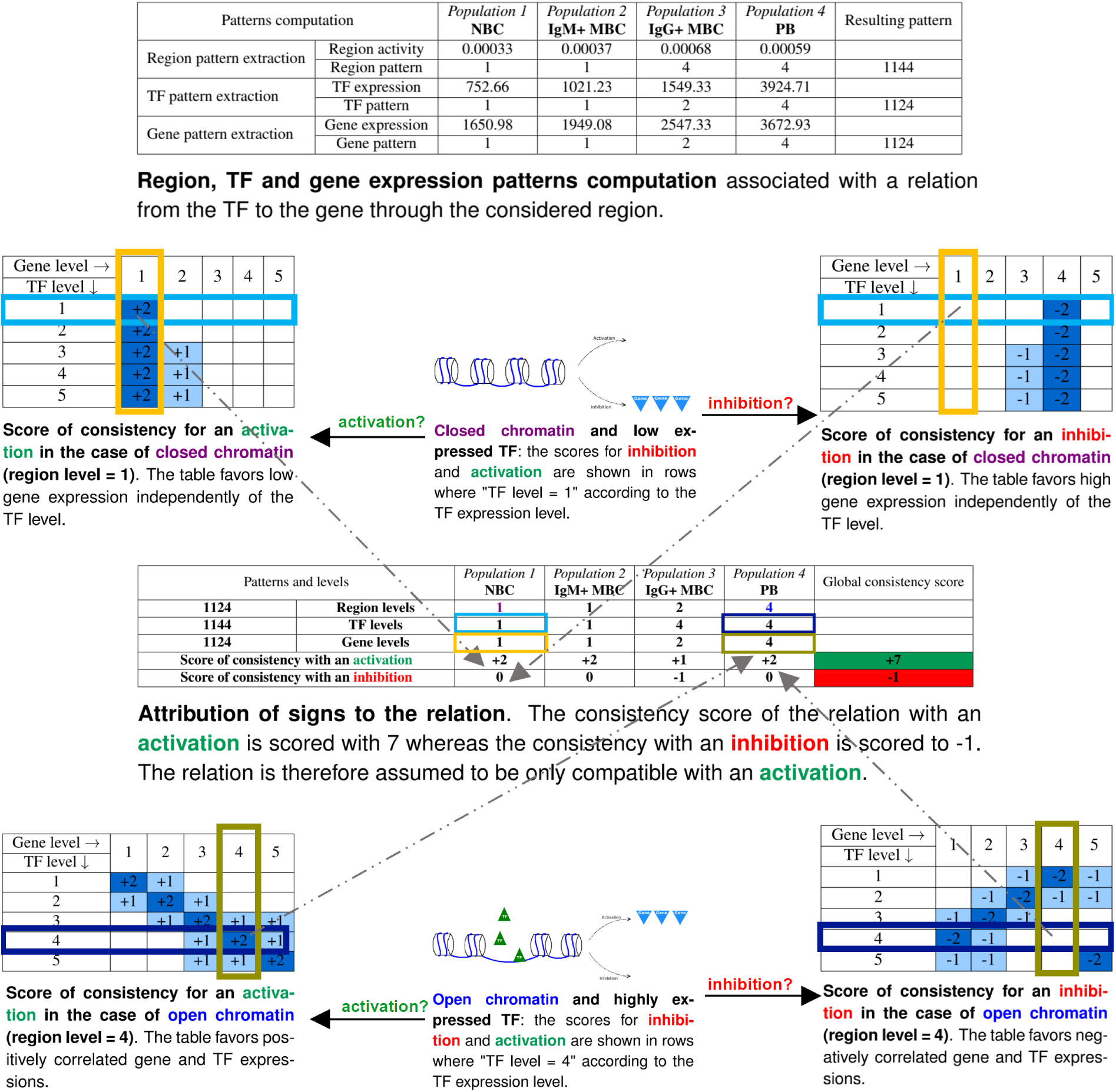
Construction of the consistency score. Top table: for each relation between a TF, a region and a gene, entities patterns are constructed as shown in Figure 5. Then for each cell population, the level of each entity pattern is used to create a consistency score. The region level and the sign of the regulation (activation in green or inhibition in red) determines which table is used among the ones presented in Supplementary Figure 3. In our example, tables for close (pattern level = 1, in purple) or highly accessible (pattern level = 4, in blue) regions are shown. Then, the intersection between TF (lines) and gene (columns) pattern levels (yellow and light blue boxes for closed regions, olive green and dark blue for open regions) determines a score, either for activation (left tables) or inhibition (right tables). This individual cell population scores are then added and submitted to a user-defined threshold to determine a global consistency score taking into account all the studied cell populations.

### 4.5. Finding key regulators with *ClassFactorY*

*Regulus* output tables can be merged to obtain unique TF-gene relations and refined by using the coverage / specificity and GO / MeSH annotation standalone external tool *ClassFactorY*.

#### 4.5.1. Coverage and specificity filters

The first filter applied by *ClassFactorY* is on coverage and specificity of TF for some gene patterns. (a) <u>Definitions and computing</u> Coverage of a TF is calculated as the proportion of genes in a specific pattern which are targets of a given TF. The coverage itself does not provide enough information: the smaller patterns, sometimes composed of 1 or 2 genes, are easily fully covered by a TF. Specificity is based on the proportion of targets genes that are from a specific pattern, for a given TF. A TF has a high specificity for a pattern if out of all its targets a significant number comes from this pattern. As for the coverage, the specificity does not yield enough information by itself: despite having a large number of its targets in a pattern, a TF may have little influence on it if the pattern is large. Both the coverage and the specificity are calculated as percentages. (b) <u>Combination of coverage and specificity</u> For a given pattern, a TF of interest is a TF which specificity and coverage are both superior to a threshold chosen by the user: mean + one standard deviation, quantiles or specific percentages. Then *ClassFactorY* outputs a list of selected TFs together with the gene patterns they potentially regulate.

#### 4.5.2. GO and MeSH annotation

To validate the TFs inferred by *Regulus*, the stand-alone module *ClassFactorY* allows for automatic queries of the GO, Uniprot and PubMed databases. A user-defined list of GO annotations is used to verify if candidates TFs are annotated by these terms. The MeSH terms are retrieved from a user-defined list of PubMed publications about the biological context and the module counts the number of citations associating the TFs to one or several of the terms. An annotation-based score is then calculated and provided as a help-decision tool for end users.

### 4.6. Datasets used for validation and testing

#### 4.6.1. Roadmap Epigenomics RNA-seq datasets

Gene inclusion in our networks is validated with *Roadmap Epigenomics* RNA-seq datasets used in (Marbach et al. 2016). RNA-seq datasets corresponding to the *FANTOM5* cell populations used for network inference are separated in three gene sets corresponding to the 10% most expressed ones, 10% less expressed ones and the 10% in the center of the expression distribution. For each category, the percentage of genes which are included in the inferred networks is reported.

#### 4.6.2. B-cells datasets

*Regulus* prediction abilities was investigated with datasets of differentiating B cells, comprising 3 replicates of RNA-seq for naive B cells (NBC), IgM secreting or IgG secreting memory B cells (IgM+ or IgG+ MBC). Data are aligned on the hg19 human reference genome and gene expressions for 26,734 genes are calculated with *featureCounts*. Raw counts files are then used for differential expression analysis with *DESeq2* (Love et al. 2014) in *R-4*.*0*.*0*. Low expressed genes are filtered out by the *R* package *HTSFilter*. Remaining genes are used for differential expression analysis by comparing each population again all the others. Variable genes are determined as those with an adjusted p-value < 0.05 and an absolute value of log2(fold_change) > 1 in at least one comparison. ATAC-seq data is obtained on the same cell types (n = 1), aligned on hg19 and accessible regions are called with MACS2 with a q-value cutoff of 0.001. An union of 58,848 regions called in at least one sample is computed with *bedtools merge* (Quinlan 2014) by taking the widest boundaries for overlapping regions. For each sample, reads overlapping each region of this union are counted for with *bedtools intersect* (Quinlan 2014) and normalized by sequencing depth and region size to compute read densities. This dataset, experimental procedures and detailed analysis settings are available on GEO (GSE136988 and GSE190458).

## DATA AVAILABILITY STATEMENT

The data underlying this article are available in the article and in its online supplementary material. *Regulus* software and source code are available on gitlab at: https://gitlab.com/teamDyliss/regulus. *ClassFactorY* software and source code are available on gitlab at: https://gitlab.com/teamDyliss/ClassFactorY.

*Regulatory Circuits* datasets and networks, FANTOM5 and GTex data used for testing and validation are available in the supplementary data of (Marbach et al. 2016) at http://www2.unil.ch/cbg/regulatorycircuits/Supplementary_data.zip and http://www2.unil.ch/cbg/regulatorycircuits/FANTOM5_individual_networks.tar. B cells datasets are available on GEO, GSE136988 (RNA-seq for PB, public) and GSE190458 (all other data, reviewer token: **wlejykemhdgpdcr**).

## ACKNOWLEDGEMENTS

We acknowledge the GenOuest bioinformatics core facility for providing the computing infrastructure (https://www.genouest.org).

## AUTHOR CONTRIBUTIONS

**ML** designed the workflow and the outline of the study. She coded the initial version. She designed the application to B cells and coded the initial version. **GC** automated and optimized the workflow. **EB** automated the validation, designed and developed *ClassFactorY* and ran the experiments on B cells. **TF** participated to the clinical expertise. **OD** participated to the design of the workflow and the study. **AS** participated to the design of the workflow and the study. **FC** contributed to the design of the workflow and the study, designed the expression patterns and designed the signed compatibility table.

## Footnotes

1 Databases were queried the 08 October 2021.

