## Supplemental Tables & Figures for "*Regulus* infers signed regulatory networks in few samples from regions and genes activities"

### Supplementary Materials

| Method name | Reference | Data | | | | Data Normalization (discretization) | | | Graph | | Implementation | Comment |
|---|---|---|---|---|---|---|---|---|---|---|---|---|
| | | Genes | Regions | TF | Other | 2 level | multi-level | continuous | Scored | Signed | | |
| **REVEAL** | (Liang et al. 1998) | a(t) | | | | x | | | | | None | |
| **RelNet** | (Butte and Kohane 1999) | a(t) | | | | x | | | x | | None | |
| **BANJO** | (Hartemink et al. 2000) | a(t) | | | | | x | | | | None | |
| **NIR** | (Gardner et al. 2003) | a(t) | | | | | x | | | | None | |
| **ARCANE** | (Margolin et al. 2006) | a(t) | | | | x | | | x | | None | |
| **TSNI** | (Bansal et al. 2006) | a(t) | | | | | x | | x | | None | Focus on 1 gene |
| **COALESCE** | (Huttenhower et al. 2009) | a(t) | | BS* | nucleosome positioning*, evolutionary conservation* | | x | | x | | C++ implementation & web interface | |
| **DISTILLER** | (Lemmens et al. 2009) | a | | BS | | | | x | x | | Integration: self mining | co-expressed genes |
| **Mix-CLR** | (Madar et al. 2010) | a(t) | | | | | x | | x | | None | |
| **TIGRESS** | (Haury et al. 2012) | a(t) | | | | | x | | x | | Matlab implementation | |
| **iRafNet** | (Petralia et al. 2015) | a(t)*, a*<br><br>Knock-down* | | BS* | interaction<br><br>protein-protein* | | x | | x | | R implementation | |
| **Regulatory Circuits** | (Marbach et al. 2016) | | a | BS | | | x | | x | | Workflow | |
| **SINCERITIES** | (Papili Gao et al. 2018) | a(t) | | | | | | x | x | x | None | |
| **PoLoBag** | (Roy et al. 2020) | a(t) | | | | | | x | x | x | None | |

Supplementary Table. 1. **Review of current network inference methods.** a = activity, a(t) time series of the activity, * optional, BS = TF binding site, None = description of the algorithm without implementation. Most methods (11/14) use time series of gene expressions as the only input data, and therefore do not take the regulatory regions activity into account. Few use information about TFs binding sites or regulatory regions (but among them, we identified *Regulatory Circuits* (Marbach et al. 2016)) and none checks whether the candidate TFs are expressed. The resulting networks may then contain relations which are not consistent with the biological situation. Most methods also produce networks with weighted edges, based on statistical or probabilistic analyses requiring large datasets acquired at several time points, which is a strong limitation to their application to human data. Indeed, many of these methods have only been tested on *Escherichia Coli* expression data and are limited to small subset of genes, raising the question of their scalability and application to human settings. Finally, we noticed that only the two most recent methods (Papili Gao et al. 2018; Roy et al. 2020) predict the activator or inhibitor role of the inferred regulations to generate signed networks, but they ignore both TF expression levels and binding site accessibility. The closest method to what we aim for is *Regulatory Circuits*, but it still shows some design and reproducibility issues, as shown in the main text. Relative to the Introduction.

| Pattern or Nb of patterns | Nb of genes in the pattern(s) | Percentage among all genes | Nb TFs in the pattern(s) | Nb of TF-genes relations targeting the pattern(s) | Percentage of all relations | Percentage of activations | TF targeting the pattern | Nb of TF passing the coverage / specificity filter | Percentage of TF passing the coverage / specificity filter | Biological interpretation |
|---|---|---|---|---|---|---|---|---|---|---|
| All: 109 | 26,734 | 100 % | 602 | 313,627 | 100 % | 50.82 %* | 275 | 121 | 44.00 % | |
| Pattern *0000* | 14,921 | 55.81 % | 327 | 0 | 0 % | 0 % | 0 | 0 | 0 % | Not expressed |
| Pattern *5555* | 3,591 | 13.43 % | 56 | 56,087 | 17.88 % | ND % | 42 | 0 | 0 % | No variations |
| Pattern *4441* | 1,418 | 5.30 % | 63 | 71,120 | 22.68 % | 64.62 % | 186 | 0 | 0 % | Decreasing expression in PB |
| Pattern *1114* | 1,244 | 4.65 % | 11 | 62,082 | 19.79 % | 38.22 % | 184 | 0 | 0 % | Increasing expression in PB |
| Patters with more than 100 genes, n = 16 | [102-557]<br>mean: 232<br>median: 173<br>sum: 3,704 | [0.38-2.08]%<br>mean: 0.87%<br>median: 0.65<br>sum: 13.86% | [0-15]<br>mean: 5<br>median: 3.5 | [650–21,026]<br>mean: 6,798<br>median: 3,172<br>sum: 108,764 | [0.21-6.70]%<br>mean: 2.17%<br>median: 1.01%<br>sum: 34.68% | [15.78-82.40]%<br>mean: 48.25%<br>median: 44.00% | [54-148]<br>mean: 117<br>median: 121 | [10-23]<br>mean: 16<br>median: 15.5 | [8.11-20.45]%<br>mean: 14.65%<br>median: 15.66% | Potentially interesting: nb genes |
| Patters with less than 100 genes, n = 92 | [[1-76]<br>mean: 21<br>median: 13<br>sum: 1,856 | [0-0.28]%<br>mean: 0.08%<br>median: 0.05%<br>sum: 6.94% | [0-4]<br>mean: 0.74<br>median: 0 | [0–1,307]<br>mean: 175<br>median: 56<br>sum: 15,574 | [0-0.42]%<br>mean: 0.06%<br>median: 0.02%<br>sum: 4.97% | [0-100]%<br>mean: 53.29%<br>median: 55.56% | [0-111]<br>mean: 35<br>median: 25 | [0-17]<br>mean: 5<br>median: 2 | [0-33.3]%<br>mean: 10.39% %<br>median: 11.71 | Difficult to interpret: low nb of genes |

Supplementary Table. 2. **Descriptive statistics on gene expression patterns and TF-gene relations obtained by applying *Regulus* to human B cell subsets.** Data were processed with a restrictive set of TF binding sites, filtered to have a strictly positive score as found in (Marbach et al. 2016) supplementary data. ND: not determined, as all consistent relations involving constantly expressed genes also involve TF and regions with constant activities; it is therefore not possible to qualify these relations as activation or inhibition. *: percentage taking into account the undetermined relations of the *5555* pattern. Relative to Figure 3 and Section 2.3.

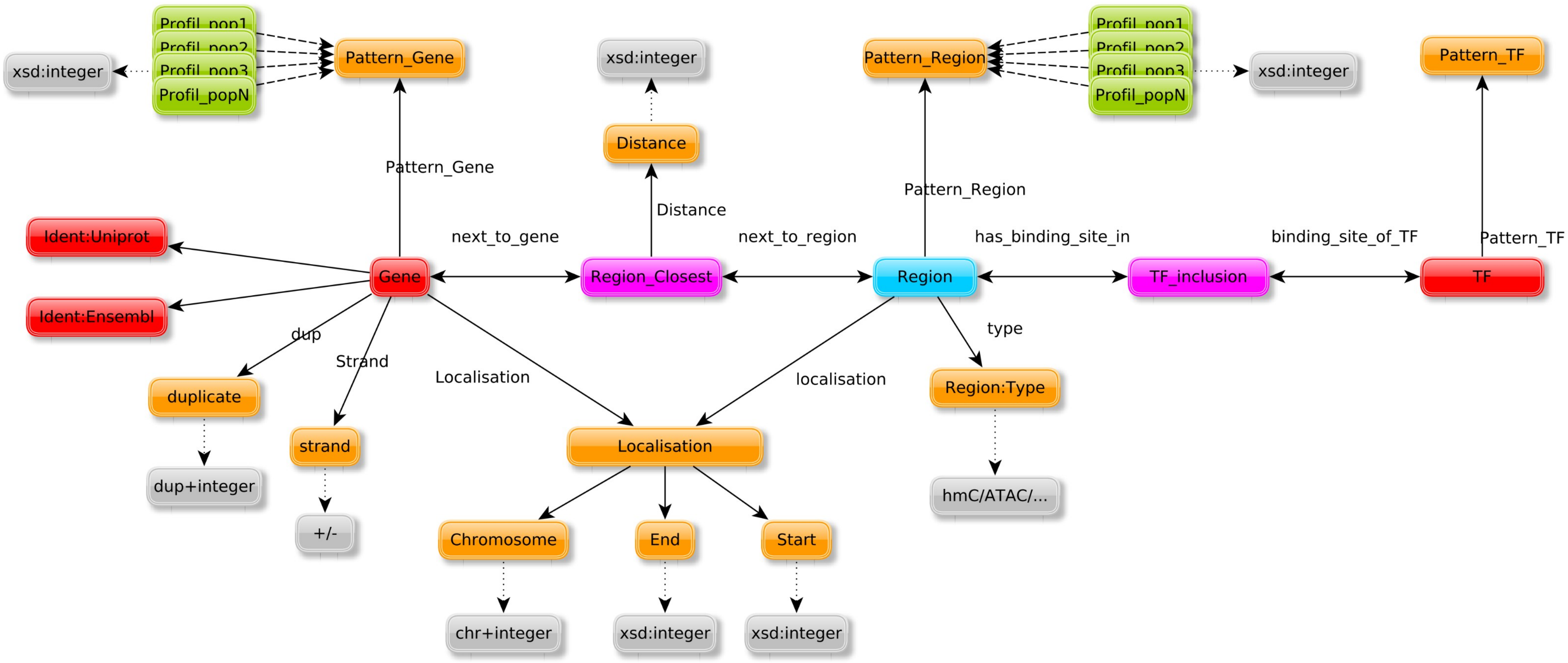


Supplementary Fig. 1. **RDF data structure model after integration of all pre-processed input data.** Example of the data structure obtained after applying Semantic Web technologies integration to B cells genomic datasets. Relative to Figure 1 and Sections 2.1.2 and 4.3

```
PREFIX : <http://www.semanticweb.org/user/ontologies/2018/1#>
PREFIX rdf: <http://www.w3.org/1999/02/22-rdf-syntax-ns#>
PREFIX rdfs: <http://www.w3.org/2000/01/rdf-schema#>

SELECT DISTINCT ?Gene ?Pattern_Gene ?Region
    ?Pattern_Region ?TF ?Pattern_TF
WHERE {
    ?Gene_uri :next_to_gene ?Region_Closest_uri .
    ?Region_Closest_uri :next_to_region ?Region_uri .
    ?TF_inclusion_uri :has_binding_site_in ?Region_uri .
    ?TF_inclusion_uri :binding_site_of_TF ?TF_uri .
    ?Gene_uri rdf:type :Gene .
    ?Gene_uri rdfs:label ?Gene .
    ?Gene_uri :PatternGene ?Pattern_GeneCategory .
    ?Pattern_GeneCategory rdfs:label ?Pattern_Gene .
    ?Region_Closest_uri rdf:type :Region_Closest .
    ?Region_Closest_uri rdfs:label ?Region_Closest .
    ?Region_Closest_uri :Distance ?Region_Closest_Distance .
    ?Region_uri rdf:type :Region .
    ?Region_uri rdfs:label ?Region .
    ?Region_uri :Pattern_Region ?Pattern_RegionCategory .
    ?Pattern_RegionCategory rdfs:label ?Pattern_Region .
    ?TF_inclusion_uri rdf:type :TF_inclusion_ATAC .
    ?TF_inclusion_uri rdfs:label ?TF_inclusion .
    ?TF_uri rdf:type :Transcription_Factor .
    ?TF_uri rdfs:label ?TF .
    ?TF_uri :PatternTF ?Pattern_TFCategory .
    ?Pattern_TFCategory rdfs:label ?Pattern_TF .
    FILTER ( ?Region_Closest_Distance < 500000 ) .

}
```

| Gene level → | 1 | 2 | 3 | 4 | 5 |
|---|---|---|---|---|---|
| TF level ↓ | | | | | |
| 1 | +2 | +1 | | | |
| 2 | +1 | +2 | +1 | | |
| 3 | | +1 | +2 | +1 | +1 |
| 4 | | | +1 | +2 | +1 |
| 5 | | | +1 | +1 | ND |

(a) Activation, Region = 5

| Gene level → | 1 | 2 | 3 | 4 | 5 |
|---|---|---|---|---|---|
| TF level ↓ | | | | | |
| 1 | | | -1 | -2 | -1 |
| 2 | | -1 | -2 | -1 | -1 |
| 3 | -1 | -2 | -1 | | |
| 4 | -2 | -1 | | | |
| 5 | -1 | -1 | | | ND |

(b) Inhibition, Region = 5

| Gene level → | 1 | 2 | 3 | 4 | 5 |
|---|---|---|---|---|---|
| TF level ↓ | | | | | |
| 1 | +2 | +1 | | | |
| 2 | +1 | +2 | +1 | | |
| 3 | | +1 | +2 | +1 | +1 |
| 4 | | | +1 | +2 | +1 |
| 5 | | | +1 | +1 | +2 |

(c) Activation, Region = 4

| Gene level → | 1 | 2 | 3 | 4 | 5 |
|---|---|---|---|---|---|
| TF level ↓ | | | | | |
| 1 | | | -1 | -2 | -1 |
| 2 | | -1 | -2 | -1 | -1 |
| 3 | -1 | -2 | -1 | | |
| 4 | -2 | -1 | | | |
| 5 | -1 | -1 | | | -2 |

(d) Inhibition, Region = 4

| Gene level → | 1 | 2 | 3 | 4 | 5 |
|---|---|---|---|---|---|
| TF level ↓ | | | | | |
| 1 | +2 | +1 | | | |
| 2 | +2 | +1 | | | +1 |
| 3 | +1 | +2 | +1 | | +1 |
| 4 | | +1 | +2 | +1 | |
| 5 | | +1 | +1 | | +2 |

(e) Activation, Region = 3

| Gene level → | 1 | 2 | 3 | 4 | 5 |
|---|---|---|---|---|---|
| TF level ↓ | | | | | |
| 1 | | | -1 | -2 | |
| 2 | | | -1 | -2 | -1 |
| 3 | | -1 | -2 | -1 | -1 |
| 4 | -1 | -2 | -1 | | |
| 5 | | -1 | -1 | | -2 |

(f) Inhibition, Region = 3

| Gene level → | 1 | 2 | 3 | 4 | 5 |
|---|---|---|---|---|---|
| TF level ↓ | | | | | |
| 1 | +2 | | | | +1 |
| 2 | +2 | +1 | | | +1 |
| 3 | +2 | +1 | | | |
| 4 | +1 | +2 | +1 | | |
| 5 | +1 | +1 | | | +1 |

(g) Activation, Region = 2

| Gene level → | 1 | 2 | 3 | 4 | 5 |
|---|---|---|---|---|---|
| TF level ↓ | | | | | |
| 1 | | | | -2 | |
| 2 | | | -1 | -2 | |
| 3 | | | -1 | -2 | -1 |
| 4 | | -1 | -2 | -1 | -1 |
| 5 | | | -1 | -1 | -1 |

(h) Inhibition, Region = 2

| Gene level → | 1 | 2 | 3 | 4 | 5 |
|---|---|---|---|---|---|
| TF level ↓ | | | | | |
| 1 | +2 | | | | |
| 2 | +2 | | | | |
| 3 | +2 | +1 | | | |
| 4 | +2 | +1 | | | |
| 5 | +2 | +1 | | | |

(i) Activation, Region = 1

| Gene level → | 1 | 2 | 3 | 4 | 5 |
|---|---|---|---|---|---|
| TF level ↓ | | | | | |
| 1 | | | | -2 | |
| 2 | | | | -2 | |
| 3 | | | -1 | -2 | |
| 4 | | | -1 | -2 | |
| 5 | | | -1 | -2 | |

(j) Inhibition, Region = 1

Supplementary Fig. 3. **Consistency scores attribution tables**, divided by relation qualification as activation (left column) or inhibition (right column) and by region pattern value from 5 (top, constant) to 1 (bottom, less accessible). In cases where all entities (TF, region and gene) have a constant activity (*5555* patterns), relations can not be qualified as activation or inhibition, as shown by the green cells labeled "ND" in the two upper tables. Relative to Figure 6 and Section 4.4.

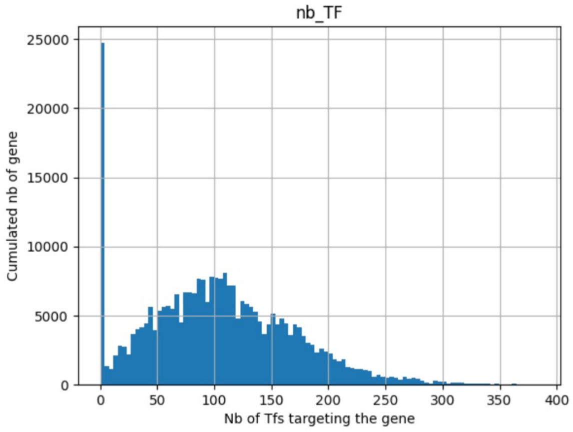


(a)

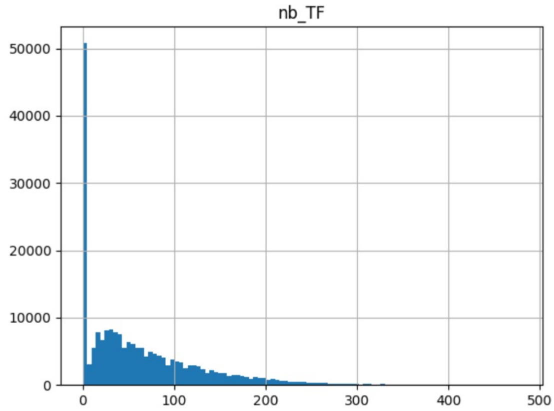


(b)

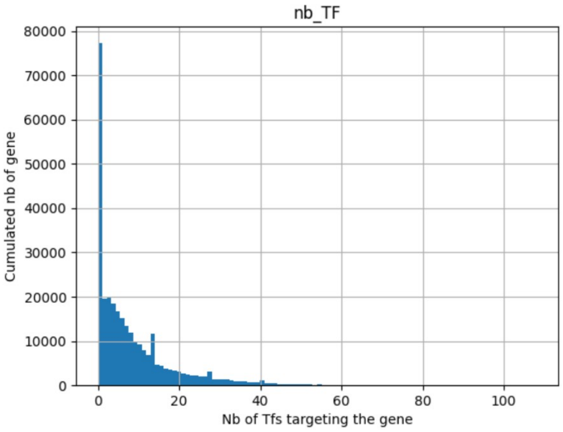


(c)

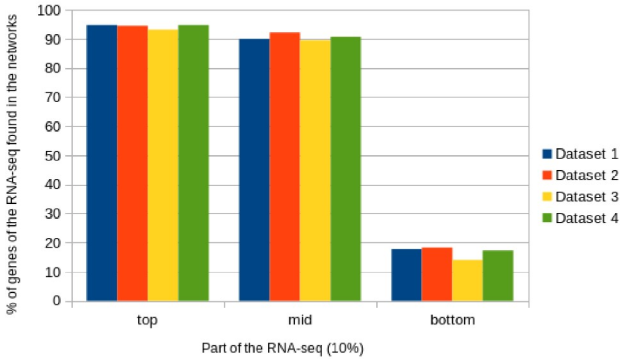


(d)

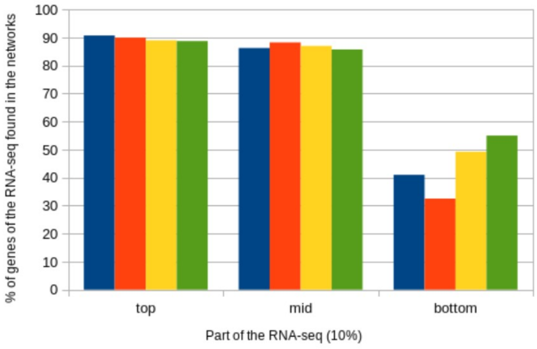


(e)

Supplementary Fig. 4. **Distribution of the number of TFs potentially regulating a gene on different network inference methods.** (a) Distribution for *Regulatory Circuits* networks, uniquely for the reduced datasets used in 2.2 and Figure 2. (b) Distribution for *Regulus* networks before applying the global consistency rules. (c) Distribution for *Regulus* final networks. All data presented here include non-expressed genes, explaining the high value for x = 0. (e-f): Percentage of genes from *Roadmap Epigenomics* RNA-seq datasets related to the cell populations found in networks inferred by *Regulatory Circuits* (e) and *Regulus* (f) according to their expression level. The datasets are the ones presented in Section 2.2 and Figure 2F. Relative to Section 2.4 and Figures 4 and 2C-D.
